# Development of a metabolic engineering technology to simultaneously suppress the expression of multiple genes in yeast and application in carotenoid production

**DOI:** 10.1101/2024.01.18.576177

**Authors:** Ryosuke Yamada, Chihiro Yamamoto, Rumi Sakaguchi, Takuya Matsumoto, Hiroyasu Ogino

## Abstract

In yeast metabolic engineering, there is a need for technologies that simultaneously suppress and regulate the expression of multiple genes and improve the production of target chemicals. In this study, we aimed to develop a novel technology that simultaneously suppresses the expression of multiple genes by combining RNA interference with global metabolic engineering technology (GMES). Furthermore, using β-carotene as the target chemical, we attempted to improve its production by using the technology. First, we developed a technology to suppress the expression of the target genes with various strengths using RNA interference. Using this technology, total carotenoid production was successfully improved by suppressing the expression of a single gene out of 10 candidate genes. Then, using this technology, RNA interference strain targeting 10 candidate genes for simultaneous suppression was constructed. The total carotenoid production of the constructed RNA interference strain was 1.7 times compared with the parental strain. In the constructed strain, the expression of eight out of the 10 candidate genes was suppressed. We developed a novel technology that can simultaneously suppress the expression of multiple genes at various intensities and succeeded in improving carotenoid production in yeast. Because this technology can suppress the expression of any gene, even essential genes, using only gene sequence information, it is considered a useful technology that can suppress the formation of by-products during the production of various target chemicals by yeast.

## Introduction

Against the backdrop of issues such as the depletion of petroleum resources and global warming, chemical production from renewable resources has attracted attention. Among the microorganisms used to produce useful chemicals, the yeast *Saccharomyces cerevisiae* is easy to cultivate on a large scale, is highly safe, and has developed a genetic recombination technology, making it a promising microorganism for industrial use^1–3^. Research is actively being conducted to improve product titers and product yield^4^.

One useful way to improve yeast product yield is to disrupt enzyme genes that contribute to by-product formation. However, out of approximately 6,000 genes in yeast, there are over 1,000 essential genes that cannot be disrupted^5^. Furthermore, in addition to essential genes, there are many genes whose disruption deteriorates growth, and cases have been reported in which the disruption of genes deteriorates growth by 90% compared to the parent strain^5^. One way to appropriately suppress the expression of genes that contribute to by-product formation without disrupting them is to replace the promoter sequence that controls gene transcription with one with a lower strength^6^.

However, using this method, it is difficult to replace the promoters of many genes simultaneously; thus, suppressing the expression of multiple genes requires considerable time and effort. Another problem is that because gene expression changes depending on the combination of the gene and promoter^7,8^, it is difficult to discover a promoter that can appropriately suppress the expression of the target gene.

To suppress gene expression using a simpler method, RNA interference, which can suppress gene expression without disrupting the genes, is effective. RNA interference is a phenomenon in which, when double-stranded RNA homologous to a portion of a target mRNA is present, the target mRNA is cleaved by proteins called Argonaute and Dicer, suppressing gene expression^9^. Although the yeast *S. cerevisiae* does not possess native RNA interference ability, studies have reported that these capabilities can be imparted by expressing Argonaute and Dicer derived from the yeast *Saccharomyces castellii*^10^. However, because a large number of genes are involved in the formation of by-products, it is difficult to know in advance which genes should be suppressed to improve the yield of the target chemical. Furthermore, there are almost no known techniques for simultaneously suppressing the expression of many enzyme-encoding genes in yeast.

In a previous study, we developed a global metabolic engineering technology (GMES) that randomly introduces multiple copies of various types of gene expression cassettes linked to genes with promoter libraries and selects optimal recombinant strains^11^. Using this technology, we succeeded in simultaneously regulating the expression of more than 10 metabolic enzyme genes and producing D-lactic acid^12^ and 2,3-butanediol^13^ with high productivity in yeast. If GMES and RNA interference technologies can be combined to optimize the expression levels of double-stranded RNA, it will be possible to develop a technology that simultaneously suppresses and regulates the expression of multiple genes and improves the product yield of target chemicals.

β-Carotene, a type of carotenoid, exhibits a high antioxidant effect, making it a valuable ingredient in cosmetics, foods, medicines, and more.^14–16^. Currently, β-carotene is mainly produced by chemical synthesis, but due to concerns about the safety of chemical synthesis, interest in production using microorganisms is increasing^14–16^. Previous research has shown that β-carotene production in *S. cerevisiae* is achieved by expressing three enzymes, phytoene synthase (crtE), lycopene cyclase (crtYB), and phytoene desaturase (crtI), derived from the yeast *Xanthophyllomyces dendrorhous*^16^. Additionally, several genes have been reported to improve carotenoid production in *S. cerevisiae* through single-gene deletions or attenuated expression (Table 1). However, there are almost no reports on improving the β-carotene yield by simultaneously suppressing the expression of multiple enzyme genes.

**Table 1.** Genes reported to increase carotenoid production by gene deletion or attenuated expression

| Gene | Relevant features | Reference |
| --- | --- | --- |
| *DPP1*<br>*LPP1*<br>*PAH1* | Encodes phosphatidate phosphatase and plays an important role in lipid synthesis | 17 |
| *ERG9* | Contributes to sterol synthesis | 18 |
| *EXG1* | Encoding glucosidase | 19 |
| *FLD1* | Controls the size and number of lipid droplets, which are storage spaces for carotenoids | 19 |
| *MNN9* | Involved in cell wall synthesis by mediating the elongation of the polysaccharide mannan skeleton | 18 |
| *ROX1* | Inhibits the expression of numerous enzymes contributing to the mevalonate pathway and ergosterol synthesis | 20 |
| *DOS2* | Unknown | 21 |
| *YJL064W* | Unknown | 21 |

In this study, we aimed to develop a novel technology that simultaneously suppresses the expression of multiple genes by combining RNA interference and GMES. Furthermore, using β-carotene as the target chemical, we attempted to improve its production by simultaneously suppressing the expression of multiple genes involved in by-product formation (Table 1) using the technology.

## Results and Discussion

### Construction of plasmids and evaluation of RNA interference performance

An episomal plasmid pEU20-Beta3 expressing *crtE*, *crtI*, and *crtYB* from *X. dendrophos*, which confers β-carotene production ability to the yeast *S. cerevisiae* was constructed (Supporting Information). In addition, because *S. cerevisiae* did not originally possess RNA interference ability, an episomal plasmid pEW-AGO-DCR that expresses Argonaute and Dicer and imparts RNA interference ability was constructed (Supporting Information). Furthermore, δ-integrative plasmid libraries pδL-ConLib* (*; DPP1, LPP1, PAH1, ERG9, EXG1, FLD1, MNN9, ROX1, DOS2, and YJL064W) that expresses double-stranded RNA to determine the target gene of RNA interference were constructed (Supporting Information).

Since pδL-ConLib* shows various double-stranded RNA expression levels depending on the number of integrated copies and the type of promoter, by introducing it together with pEW-AGO-DCR that expresses Argonaute and Dicer, it is expected that the gene of interest can be suppressed with various strengths. Therefore, we conducted an experiment to suppress GFP expression using an RNA interference system (Supplementary Figs. 2 and 3). As a result, it was confirmed that the expression of the target gene could be stably suppressed over a long period at various strengths.

### Evaluation of total carotenoid production in yeast strains targeting a single gene with RNA interference

β-Carotene-producing yeast YPH499/Beta3 was constructed by introducing the β-carotene-producing plasmid pEU20-Beta3 into a yeast strain YPH499. Plasmid pEW-AGO-DCR, which confers RNA interference ability, and plasmid pδL-ConLib*, which expresses double-stranded RNA for the target gene of RNA interference, were introduced into YPH499/Beta3. The constructed yeast strains in which the expression of a single gene was suppressed were named YPH499/Beta3/i-* (*; DPP1, LPP1, PAH1, ERG9, EXG1, FLD1, MNN9, ROX1, DOS2, and YJL064W).

The constructed yeast strains YPH499/Beta3/i-* were cultured in a synthetic complete drop-out (SCD) medium for 96 h, and total carotenoid production was measured (Fig. 1). All yeast strains subjected to RNA interference produced higher amounts of carotenoids than the parental strain YPH499/Beta3. The highest carotenoid production was observed in the YPH499/Beta3/i-ERG9 strain (17.7 mg/L), which targeted *ERG9* for RNA interference, and its value was 1.3-fold higher than the parental strain YPH499/Beta3 (13.1 mg/L).

**Fig. 1.**
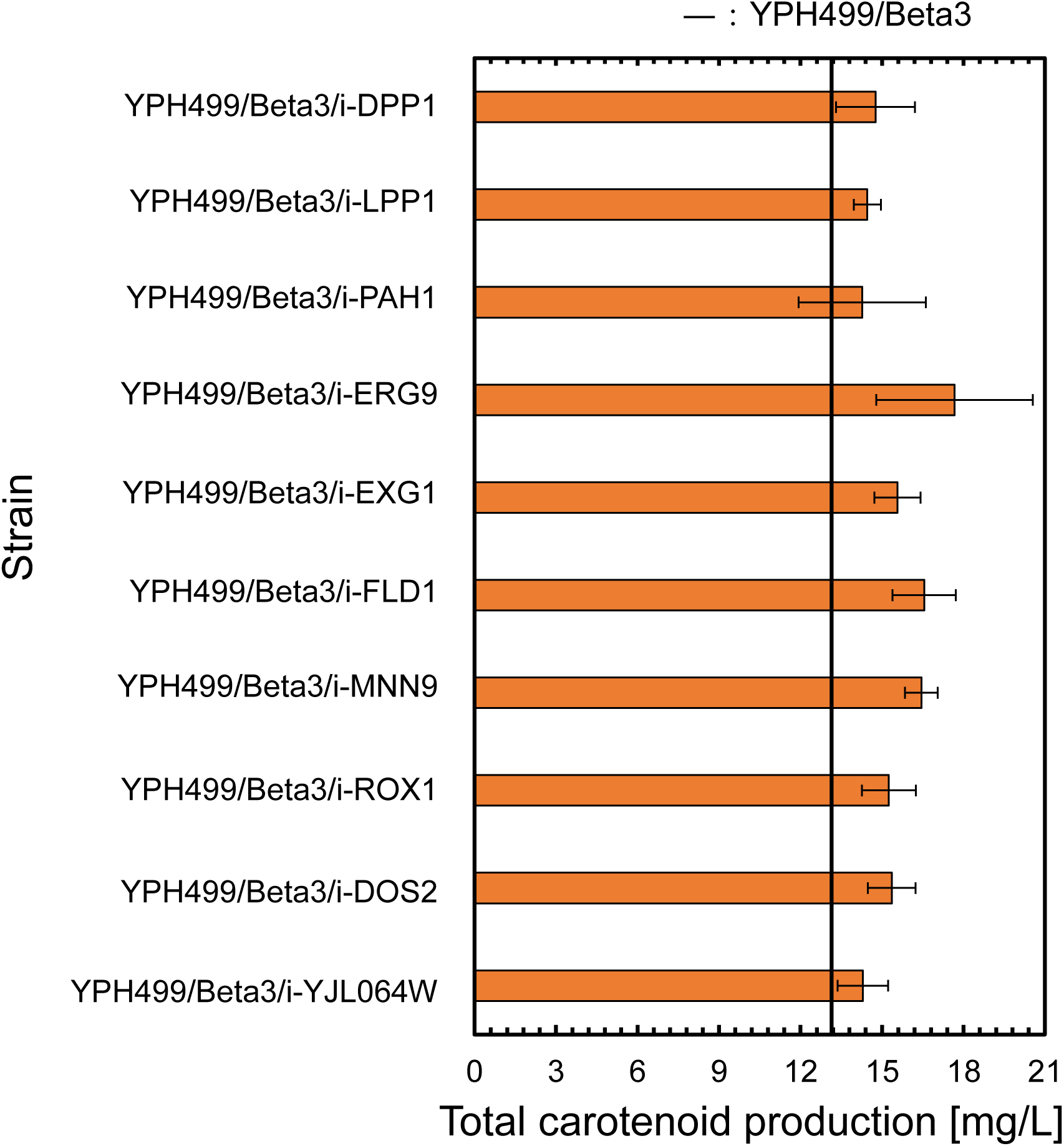
Effect of suppressing the expression of a single gene on total carotenoid production. Data represent the average of three independent experiments, and the error bar shows the standard deviation. The solid line indicates the value for the parent strain YPH499/Beta3.

*ERG9* is a gene that encodes an enzyme that synthesizes squalene from farnesyl pyrophosphate, a precursor in β-carotene biosynthesis^18^. That is, ERG9 contributes to the generation of by-products in β-carotene biosynthesis. However, *ERG9* is an essential gene that cannot be disrupted. A previous study reported that replacing the promoter of *ERG9* in yeast *S. cerevisiae* with a weak promoter and suppressing its expression reduced the production of the by-products squalene and ergosterol and improved β-carotene production by 1.3-fold compared to the parent strain^22^. Similarly, in the present study, RNA interference in YPH499/Beta3/i-ERG 9 may have suppressed *ERG9* expression, increased the metabolic flux to the β-carotene biosynthetic pathway, and increased carotenoid production.

### Confirmation of RNA interference ability in the β-carotene-producing yeast strain

To confirm that the increased carotenoid production capacity of the RNA interference strain was due to RNA interference, YPH499/Beta3/AD and YPH499/Beta3/dsR were constructed by introducing only one of the plasmids pEW-AGO-DCR or pδL-ConLibERG9, which are Argonaute and Dicer or double-stranded RNA expressing plasmid, into YPH499/Beta3.

Total carotenoid production was evaluated after culturing YPH499/Beta3, YPH499/Beta3/AD, YPH499/Beta3/dsR, and YPH499/Beta3/i-ERG 9 cells in the SCD medium for 96 h (Fig. 2). Carotenoid production by YPH499/Beta3/AD, which expressed only Argonaute and Dicer, was slightly lower than that of the parent strain. Furthermore, carotenoid production by YPH499/Beta3/dsR expressing only double-stranded RNA was comparable to that of the parent strain YPH499/Beta3. In contrast, YPH499/Beta3/i-ERG9, which expressed Argonaute, Dicer, and double-stranded RNA, showed improved carotenoid production compared with the parent strain. Thus, it was confirmed that the expression of Argonaute, Dicer, and double-stranded RNAs is necessary for RNA interference in yeast.

**Fig. 2.**
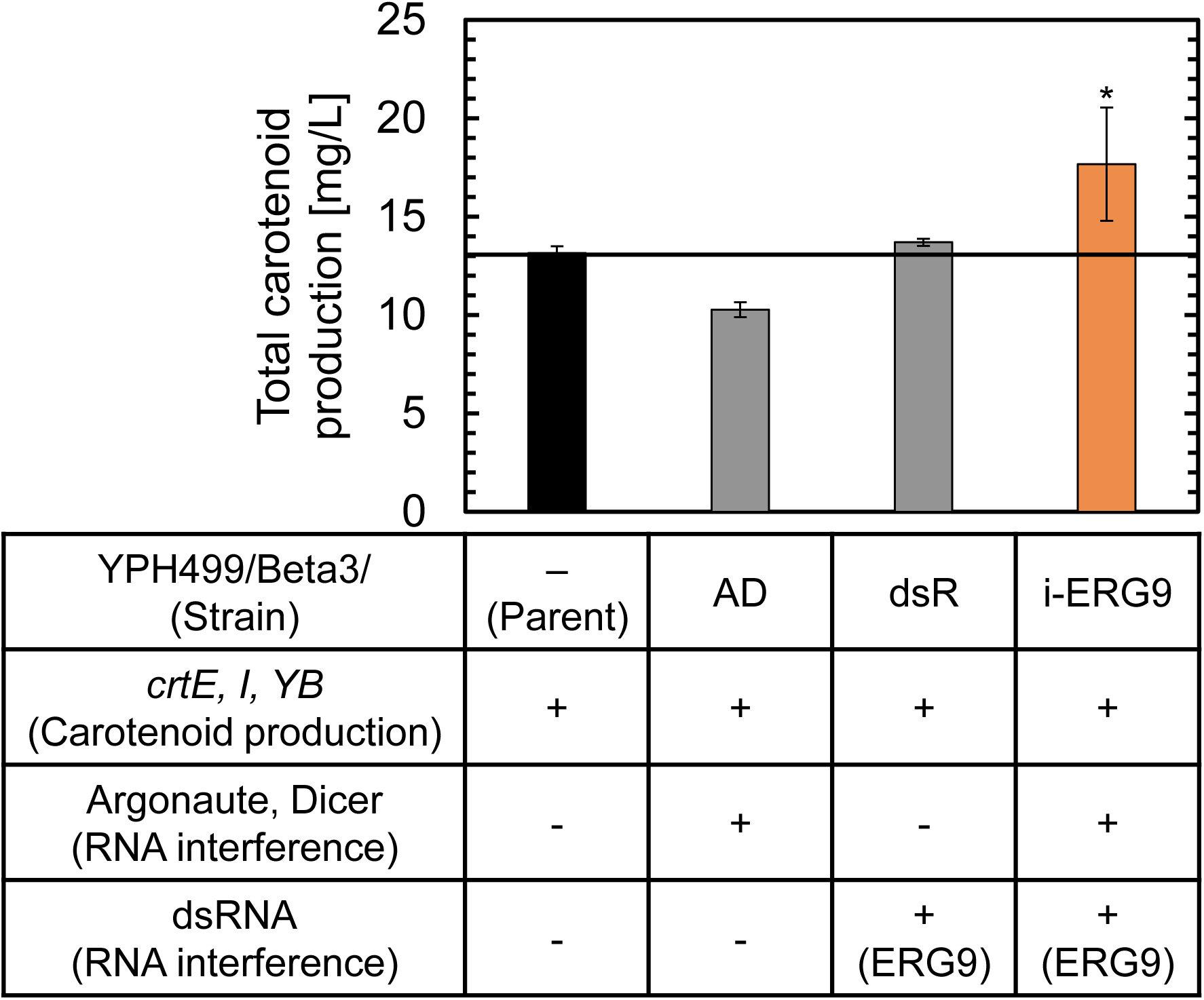
Confirmation of RNA interference ability in the β-carotene-producing strain. Data represent the average of three independent experiments, and the error bar shows the standard deviation. The solid line indicates the value for the parent strain YPH499/Beta3. * indicates a significant difference (Student’s t-test, p = 0.92).

### Evaluation of total carotenoid production in yeast strains that simultaneously targets 10 different genes for RNA interference

Using the β-carotene-producing yeast YPH499/Beta3 as the parent strain, a yeast library in which the expression of up to 10 genes was simultaneously suppressed by simultaneously introducing plasmid pEW-AGO-DCR, which confers RNA interference ability, and 10 plasmids pδL-ConLib*, which express double-stranded RNA against target genes for RNA interference was constructed. From the constructed yeast library, 181 strains with a dark red-orange color on a solid medium were isolated and named YPH499/Beta3/i-10_* (*;1-181). The isolated yeast strains YPH499/Beta3/i-10_* were cultured in SCD medium for 96 h, and total carotenoid production was determined (Fig. 3). Of the 181 strains evaluated, 48 showed higher carotenoid production than the parent strain YPH499/Beta3. The value for strain YPH499/Beta3/i-10_176, which showed the highest carotenoid production, was 21.9 mg/L. This value was 1.7-fold higher than that of the parent strain YPH499/Beta3 (13.1 mg/L) and 1.2-fold higher than that of YPH499/Beta3/i-ERG9 (17.7 mg/L), in which only *ERG9* expression was suppressed.

**Fig. 3.**
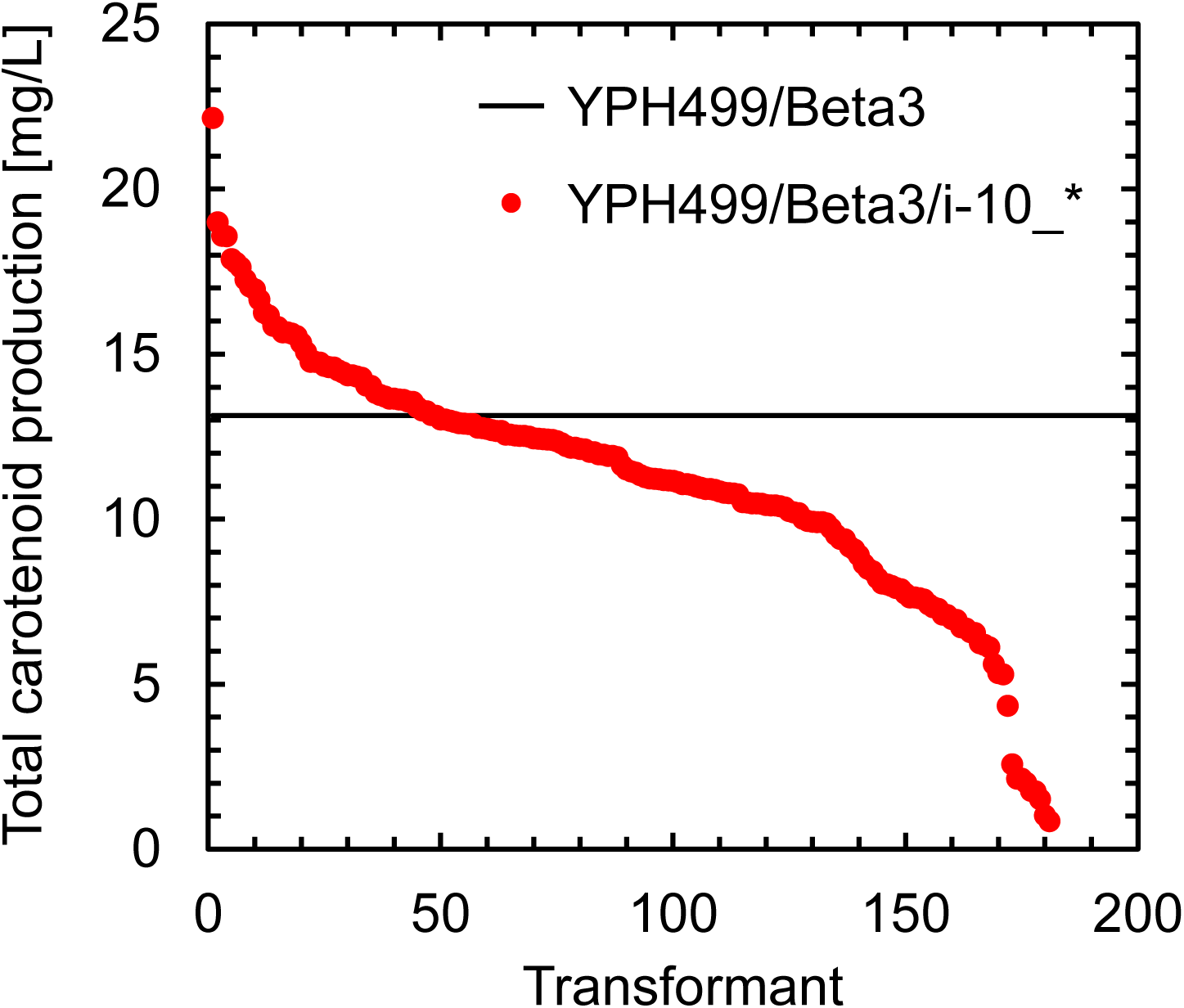
Total carotenoid production of yeast library targeting 10 genes simultaneously with RNA interference. The data were obtained from a single experiment. The solid line indicates the value for the parent strain YPH499/Beta3.

In conventional yeast gene deletion methods, only one gene is deleted in a single operation^23^. Alternatively, previous studies have reported that genome engineering technologies called clustered regularly interspaced short palindromic repeats (CRISPR)-PCR-mediated chromosomal deletion (PCD) (CRISPR-PCD) and PCR-mediated chromosomal replacement (CRISPR-PCRep) can replace three chromosomal regions in yeast at once^24^. It has also been reported that quadruple deletion of genes and simultaneous transcriptional regulation of the three genes are possible using CRISPR and the endoribonuclease Csy4^25^. However, to the best of our knowledge, no technology has been reported that can simultaneously suppress the expression of approximately 10 genes located at different locations in the genome and regulate the degree of suppression, as in this study. Therefore, this novel technology, which can simultaneously suppress the expression of 10 different genes in a single operation, is considered useful and can be applied to various situations in metabolic engineering.

### Time course of carotenoid production in RNA interference strain

YPH499/Beta3/i-10_176, which showed the highest carotenoid production by simultaneously targeting 10 different genes for RNA interference, and its parent strain YPH499/Beta3 were cultured in SCD medium, and their carotenoid production was evaluated (Fig. 4).

**Fig. 4.**
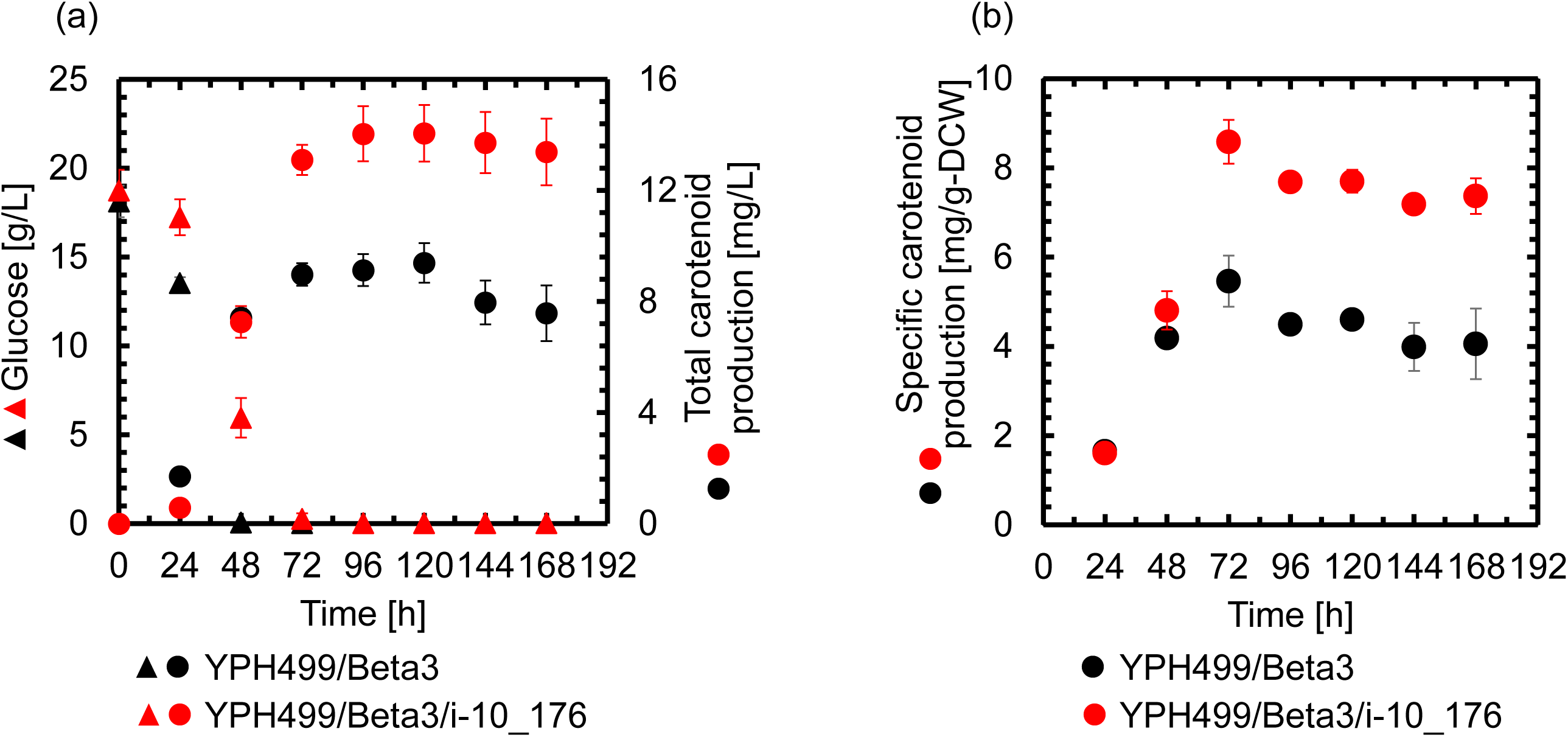
Time course of (a) glucose concentration and total carotenoid production and (b) specific carotenoid production in the RNA interference strain and its parent strain. Data are the average of three independent experiments and the error bar shows the standard deviation.

The parent strain YPH499/Beta3 completely consumed glucose after 48 h of cultivation and produced up to 9.4 mg/L of carotenoids after 120 h. The RNA interference strain YPH499/Beta3/i-10_176 completely consumed glucose after 72 h of cultivation and produced up to 14.1 mg/L of carotenoids after 120 h of culture. This value was 1.5 times higher than that of the parental strain YPH499/Beta3. The parent strain YPH499/Beta3 showed the highest specific production of 5.46 mg/g-dry cell weight (DCW) after 72 h of cultivation. In contrast, YPH499/Beta3/i-10_176 showed a maximum specific production of 8.58 mg/g-DCW at 72 h of cultivation, a 1.6-fold improvement over YPH499/Beta3.

In this study, RNA interference targeting 10 different genes improved carotenoid production, achieving a maximum carotenoid production of 8.58 mg/g-DCW per dry cell. However, in a previous study, a β-carotene production of 25.5 mg/g-DCW was achieved by culturing an engineered strain of yeast *S. cerevisiae* in bioreactors under optimal culture conditions^26^. Therefore, there is significant room for improvement in the carotenoid production of YPH499/Beta3/i-10_176 obtained in this study. Overexpression of enzyme genes that contribute to the production of target chemicals and disruption or attenuation of enzyme genes that contribute to the production of by-products are both effective in improving product yields. In the present study, we suppressed the expression of genes that inhibit β-carotene production to improve carotenoid production in yeast. Previous studies have reported that overexpression of several genes in the mevalonate pathway, as well as *crtE*, *crtYB*, and *crtI*, can increase carotenoid production^27,28^. Carotenoid production was greatly enhanced by simultaneously suppressing the expression of *ERG9* and overexpressing *HMG1*, which encodes a rate-limiting enzyme in the mevalonate pathway^18^. In the future, it is expected that the overexpression of enzyme genes that contribute to carotenoid production will be combined with the suppression of the simultaneous expression of multiple genes involved in the formation of by-products using the developed technology, leading to the development of strains with higher carotenoid production.

### Relative transcript levels of genes targeted for RNA interference

The relative transcription levels of YPH499/Beta3/i-10_176, in which 10 types of genes were simultaneously targeted by RNA interference, relative to the parent strain YPH499/Beta3, were evaluated (Fig. 5).

**Fig. 5.**
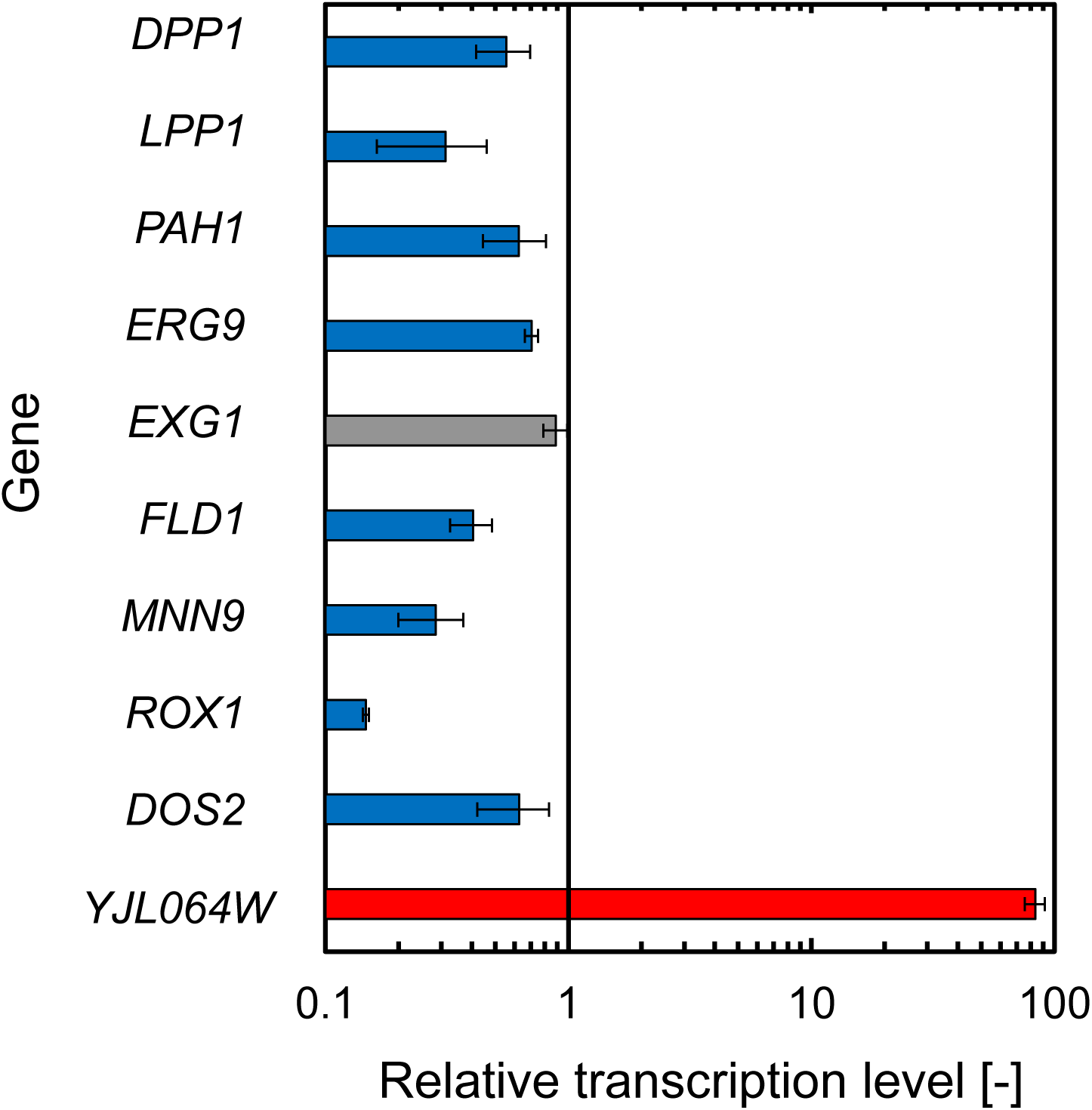
Relative transcript levels of genes targeted for RNA interference in the RNA interference strain. Data represent the average of three independent experiments, and the error bar shows the standard deviation. Red and blue bars represent significant increases and decreases, respectively, relative to the parent strain YPH499/Beta3.

In YPH499/Beta3/i-10_176, the transcript levels of eight genes, except *EXG1* and *YJL064W*, significantly decreased at various rates, ranging from 0.15 to 0.71 times. These results revealed that the expression of eight types of genes was suppressed to various degrees in YPH499/Beta3/i-10_176 compared to that in the parent strain YPH499/Beta3. In contrast, the transcription level of YJL064W increased 83.6 times.

Interestingly, in the YPH499/Beta3/i-10_176 strain, the expression of eight genes decreased owing to RNA interference, but the transcription level of *YJL064W* increased. Regulation of gene expression levels involves the expression levels of various other genes and intracellular metabolite concentrations^29^; possible causes for the increased expression of *YJL064W* include changes in other gene expression levels and associated changes in intracellular metabolic fluxes and metabolite concentrations.

Previous studies have reported that although disruption of each gene alone does not improve carotenoid production, simultaneous disruption of multiple genes improves carotenoid production in yeast^30^. The novel technology established in this study, which combines RNA interference and GMES, can target any gene, even essential genes, as long as the gene sequence is known, and multiple genes can be targeted for interference at the same time. Therefore, this technology can be applied to various yeast genes to identify combinations of genes whose expression suppression contributes to the improvement of product yield in the production of various chemicals, which has not yet been clarified.

## Conclusions

In this study, we developed a novel technology that can simultaneously suppress the expression of multiple genes at various intensities by combining RNA interference and GMES and succeeded in improving carotenoid production in yeast. Because this novel technology can suppress the expression of any gene, even essential genes, using only gene sequence information, it is considered a useful technology that can suppress the formation of by-products during the production of various target chemicals by yeast.

## Materials and Methods

### Strains and medium

*Escherichia coli* strain HST08 (Takara Bio, Otsu, Japan) was used as the host for recombinant DNA manipulation. Recombinant *E. coli* cells were cultured in Luria–Bertani (LB) medium made with 20 g/L LB broth powder (Nacalai Tesque, Kyoto, Japan) supplemented with 100 μg/mL ampicillin sodium salt.

*S. cerevisiae* strains were cultivated in SCD medium (6.7 g/L yeast nitrogen base without amino acids (Formedium, Norfolk, UK), 2.0 g/L synthetic complete mixture (Formedium), and 20 g/L glucose) supplemented with the appropriate amino acids and nucleic acids. For the solid medium, 20 g/L of agar was added to the medium.

### Yeast cultivation

Microplate cultivation was performed in 1.2 mL of SCD medium for 72 h using a 2 mL 96-deep well plate equipped with a gas-permeable seal and a rotary plate shaker set at 30°C and 1500 rpm. Cultivation was initiated by inoculation (5% vol/vol) of a preculture grown on a microplate containing SCD medium for 16 h at 30°C and 1500 rpm.

Flask cultivation was performed in 200 mL of SCD medium using a rotary shaker operated at 30°C and 150 rpm. Cultivation was initiated by inoculation (initial OD_600_: 0.05) of a preculture grown for 24 h at 30°C in a test tube containing SCD medium at 150 rpm.

### Plasmid construction

β-Carotene producing plasmid pEU20-Beta3, Argonaute and Dicer expressing plasmid pEW-AGO-DCR, and double-stranded RNA expressing plasmid pδL-ConLib* were constructed (Supporting Information). The constructed plasmid was introduced into *S. cerevisiae* YPH499 (NBRC 10505; NITE Biological Resource Center) using a previously described lithium acetate method^31^.

### Real-time PCR

Total RNA was isolated from yeast cells cultivated in SCD medium for 72 h at 30°C, and cDNA was synthesized as previously described^32^. Gene transcription levels were quantified by real-time PCR, as described previously^32^, using cDNA as the template and pyruvate dehydrogenase (*PDA1*) as the housekeeping gene^32^. The primers used for real-time PCR are listed in Supplementary Table 1.

### Analyses

Total carotenoid^27^ and glucose^33^ concentrations were determined as previously described. Yeast DCW was measured using the previously described method^34^. Specific carotenoid production was calculated using the following equation (1).

## Author Contributions

RY: Conceptualization, Writing-Original Draft, Writing-Review & Editing, Supervision, Funding acquisition; CY: Investigation, Writing-Original Draft; RS: Investigation; TM and HO: Supervision

## Acknowledgments

We would like to thank Editage for the English language editing.

## Notes

The authors declare no competing financial interest.

## References

1 Borodina, I.; Nielsen, J. Advances in Metabolic Engineering of Yeast *Saccharomyces cerevisiae* for Production of Chemicals. Biotechnol. J. 2014, 9, 609–620.

2 Shi, B.; Ma, T.; Ye, Z.; Li, X.; Huang, Y.; Zhou, Z.; Ding, Y.; Deng, Z.; Liu, T. Systematic Metabolic Engineering of *Saccharomyces cerevisiae* for Lycopene Overproduction. J. Agric. Food Chem. 2019, 67, 11148–11157.

3 Li, S.; Zhang, Q.; Wang, J.; Liu, Y.; Zhao, Y.; Deng, Y. Recent Progress in Metabolic Engineering of *Saccharomyces cerevisiae* for the Production of Malonyl-CoA Derivatives. J. Biotechnol. 2021, 325, 83–90.

4 Lian, J.; Mishra, S.; Zhao, H. Recent Advances in Metabolic Engineering of *Saccharomyces cerevisiae*: New Tools and Their Applications. Metab. Eng. 2018, 50, 85–108.

5 Giaever, G.; Chu, A. M.; Ni, L.; Connelly, C.; Riles, L.; Véronneau, S.; Dow, S.; Lucau-Danila, A.; Anderson, K.; André, B.;, et al. Functional Profiling of the *Saccharomyces cerevisiae* Genome. Nature. 2002, 418, 387–391.

6 Shiba, Y.; Paradise, E. M.; Kirby, J.; Ro, D. K.; Keasling, J. D. Engineering of the Pyruvate Dehydrogenase Bypass in *Saccharomyces cerevisiae* for High-Level Production of Isoprenoids. Metab. Eng. 2007, 9, 160–168.

7 Da Silva, N. A.; Srikrishnan, S. Introduction and Expression of Genes for Metabolic Engineering Applications in *Saccharomyces cerevisiae*. FEMS Yeast Res. 2012, 12, 197–214.

8 Yamada, R.; Kimoto, Y.; Ogino, H. Combinatorial Library Strategy for Strong Overexpression of the Lipase From *Geobacillus thermocatenulatus* on the Cell Surface of Yeast *Pichia pastoris*. Biochem. Eng. J. 2016, 113, 7–11.

9 Agrawal, N.; Dasaradhi, P. V. N.; Mohmmed, A.; Malhotra, P.; Bhatnagar, R. K.; Mukherjee, S. K. R. N. A. RNA Interference: Biology, Mechanism, and Applications. Microbiol. Mol. Biol. Rev. 2003, 67, 657–685.

10 Drinnenberg, I. A.; Weinberg, D. E.; Xie, K. T.; Mower, J. P.; Wolfe, K. H.; Fink, G. R.; Bartel, D. P. RNAi in Budding Yeast. Science. 2009, 326, 544–550.

11 Yamada, R.; Wakita, K.; Ogino, H. Global Metabolic Engineering of Glycolytic Pathway via Multicopy Integration in *Saccharomyces cerevisiae*. ACS Synth. Biol. 2017, 6, 659–666.

12 Yamada, R.; Wakita, K.; Mitsui, R.; Ogino, H. Enhanced D-Lactic Acid Production by Recombinant *Saccharomyces cerevisiae* Following Optimization of the Global Metabolic Pathway. Biotechnol. Bioeng. 2017, 114, 2075–2084.

13 Yamada, R.; Wakita, K.; Mitsui, R.; Nishikawa, R.; Ogino, H. Efficient Production of 2,3-Butanediol by Recombinant *Saccharomyces cerevisiae* Through Modulation of Gene Expression by Cocktail Delta-Integration. Bioresour. Technol. 2017, 245, 1558–1566.

14 Li, J.; Shen, J.; Sun, Z.; Li, J.; Li, C.; Li, X.; Zhang, Y. Discovery of Several Novel Targets That Enhance β-Carotene Production in *Saccharomyces cerevisiae*. Front. Microbiol. 2017, 8, 1116.

15 Mata-Gómez, L. C.; Montañez, J. C.; Méndez-Zavala, A.; Aguilar, C. N. Biotechnological Production of Carotenoids by Yeasts: An Overview. Microb. Cell Fact 2014, 13, 12.

16 Verwaal, R.; Wang, J.; Meijnen, J. P.; Visser, H.; Sandmann, G.; van den Berg, J. A.; van Ooyen, A. J. High-Level Production of Beta-Carotene in *Saccharomyces cerevisiae* by Successive Transformation With Carotenogenic Genes From *Xanthophyllomyces dendrorhous*. Appl. Environ. Microbiol. 2007, 73, 4342–4350.

17 Zhao, Y.; Zhang, Y.; Nielsen, J.; Liu, Z. Production of β-Carotene in *Saccharomyces cerevisiae* Through Altering Yeast Lipid Metabolism. Biotechnol. Bioeng. 2021, 118, 2043–2052.

18 Lian, J.; HamediRad, M.; Hu, S.; Zhao, H. Combinatorial Metabolic Engineering Using an Orthogonal Tri-functional CRISPR System. Nat. Commun. 2017, 8, 1688.

19 Ma, T.; Shi, B.; Ye, Z.; Li, X.; Liu, M.; Chen, Y.; Xia, J.; Nielsen, J.; Deng, Z.; Liu, T. Lipid Engineering Combined With Systematic Metabolic Engineering of *Saccharomyces cerevisiae* for High-Yield Production of Lycopene. Metab. Eng. 2019, 52, 134–142.

20 Özaydın, B.; Burd, H.; Lee, T. S.; Keasling, J. D. Carotenoid-Based Phenotypic Screen of the Yeast Deletion Collection Reveals New Genes With Roles in Isoprenoid Production. Metab. Eng. 2013, 15, 174–183.

21 Chen, Y.; Xiao, W.; Wang, Y.; Liu, H.; Li, X.; Yuan, Y. Lycopene Overproduction in *Saccharomyces cerevisiae* Through Combining Pathway Engineering With Host Engineering. Microb. Cell Fact 2016, 15, 113.

22 Bu, X.; Lin, J. Y.; Duan, C. Q.; Koffas, M. A. G.; Yan, G. L. Dual Regulation of Lipid Droplet-Triacylglycerol Metabolism and *ERG9* Expression for Improved β-Carotene Production in *Saccharomyces cerevisiae*. Microb. Cell Fact. 2022, 21, 3.

23 Akada, R.; Kitagawa, T.; Kaneko, S.; Toyonaga, D.; Ito, S.; Kakihara, Y.; Hoshida, H.; Morimura, S.; Kondo, A.; Kida, K. PCR-Mediated Seamless Gene Deletion and Marker Recycling in *Saccharomyces cerevisiae*. Yeast. 2006, 23, 399–405.

24 Easmin, F.; Sasano, Y.; Kimura, S.; Hassan, N.; Ekino, K.; Taguchi, H.; Harashima, S. CRISPR-PCD and CRISPR-PCRep: Two Novel Technologies for Simultaneous Multiple Segmental Chromosomal Deletion/Replacement in *Saccharomyces cerevisiae*. J. Biosci. Bioeng. 2020, 129, 129–139.

25 Ferreira, R.; Skrekas, C.; Nielsen, J.; David, F. Multiplexed CRISPR/Cas9 Genome Editing and Gene Regulation Using Csy4 in *Saccharomyces cerevisiae*. ACS Synth. Biol. 2018, 7, 10–15.

26 Olson, M. L.; Johnson, J.; Carswell, W. F.; Reyes, L. H.; Senger, R. S.; Kao, K. C. Characterization of an Evolved Carotenoids Hyper-producer of *Saccharomyces cerevisiae* Through Bioreactor Parameter Optimization and Raman Spectroscopy. J. Ind. Microbiol. Biotechnol. 2016, 43, 1355–1363.

27 Shimazaki, S.; Yamada, R.; Yamamoto, Y.; Matsumoto, T.; Ogino, H. Building a Machine-Learning Model to Predict Optimal Mevalonate Pathway Gene Expression Levels for Efficient Production of a Carotenoid in Yeast. Biotechnol. J. 2023, e2300285.

28 Yamada, R.; Ando, K.; Sakaguchi, R.; Matsumoto, T.; Ogino, H. Simultaneous Point and Structural Mutations in Engineered Yeast *Saccharomyces cerevisiae* Improve Carotenoid Production, Research Square, 2023, doi.org/10.21203/rs.3.rs-3623691/v1.

29 Hahn, S.; Young, E. T. Transcriptional Regulation in *Saccharomyces cerevisiae*: Transcription Factor Regulation and Function, Mechanisms of Initiation, and Roles of Activators and coactivators. Genetics. 2011, 189, 705–736.

30 Trikka, F. A.; Nikolaidis, A.; Athanasakoglou, A.; Andreadelli, A.; Ignea, C.; Kotta, K.; Argiriou, A.; Kampranis, S. C.; Makris, A. M. Iterative Carotenogenic Screens Identify Combinations of Yeast Gene Deletions That Enhance Sclareol Production. Microb. Cell Fact. 2015, 14, 60.

31 Chen, D. C.; Yang, B. C.; Kuo, T. T. One-Step Transformation of Yeast in Stationary Phase. Curr. Genet. 1992, 21, 83–84.

32 Mitsui, R.; Nishikawa, R.; Yamada, R.; Matsumoto, T.; Ogino, H. Construction of Yeast Producing Patchoulol by Global Metabolic Engineering Strategy. Biotechnol. Bioeng. 2020, 117, 1348–1356.

33 Yamada, R.; Kumata, Y.; Mitsui, R.; Matsumoto, T.; Ogino, H. Improvement of Lactic Acid Tolerance by Cocktail δ-Integration Strategy and Identification of the Transcription Factor *PDR3* Responsible for Lactic Acid Tolerance in Yeast *Saccharomyces cerevisiae*. World J. Microbiol. Biotechnol. 2021, 37, 19.

34 Yamada, R.; Kashihara, T.; Ogino, H. Improvement of Lipid Production by the Oleaginous Yeast *Rhodosporidium toruloides* Through UV Mutagenesis. World J. Microbiol. Biotechnol. 2017, 33, 99.

