## Supporting Information for "Development of a metabolic engineering technology to simultaneously suppress the expression of multiple genes in yeast and application in carotenoid production"

**Plasmid construction**

All primers used for plasmid construction are listed in Supplementary Table 1. Episomal plasmid pEU20-Beta3 (Supplementary Fig. 1a) for β-carotene production was constructed using the following procedure. Briefly, a DNA fragment containing selective marker URA3 was obtained via PCR with primers URA3P_TP20-AscI(F)_Lig and pE-AscI(R)_Lig and using pEUPG (Yamada et al. 2017) as a template. The resulting DNA fragment was digested by restriction enzyme AscI and self-ligated. The resulting plasmid was named pEU20PG. Gene fragments A1–A7 were obtained via PCR with the template DNA and primers (Supplementary Table 2). The resulting DNA fragments were ligated in yeast cells (Ma et al. 1987). The resulting plasmid was named pEU20-Beta3.

Episomal plasmid pEW-AGO-DCR (Supplementary Fig. 1b) for expressing Argonaute and Dicer from *Saccharomyces castellii* was constructed using the following procedure. Briefly, gene fragments B1–B5 were obtained via PCR with the template DNA and primers (Supplementary Table 2). The resulting DNA fragments were ligated by using overlap extension PCR and NEBuilder HiFi DNA Assembly Master Mix (New England Biolabs Japan, Tokyo, Japan). The resulting plasmid was named pRS404-AGO-DCR. A gene fragments B6 was obtained via PCR with the template DNA and primers (Supplementary Table 2). The resulting DNA fragment was inserted into the XhoI site of pRS404-AGO-DCR by using NEBuilder HiFi DNA Assembly Master Mix. The resulting plasmid was named pEW-AGO-DCR.

δ-Integrative plasmid library pδL-ConLib* for expressing double-stranded RNA was constructed using the following procedure. Briefly, gene fragments C1 and C2 were obtained via PCR with the template DNA and primers (Supplementary Table 2). The resulting DNA fragments were inserted into the NotI/PstI site of pδL (ref) by using NEBuilder HiFi DNA Assembly Master Mix. The resulting plasmid library was named pδL-ConLib. Gene fragments C3–C13 containing the internal sequences of ten genes targeted by RNA interference (Supplementary Table 3) were obtained via PCR with the template DNA and primers (Supplementary Table 2). The resulting DNA fragments were inserted into the SbfI site of pδL-ConLib by using NEBuilder HiFi DNA Assembly Master Mix. The resulting plasmid libraries were named pδL-ConLib* (*; DPP1, LPP1, PAH1, ERG9, EXG1, FLD1, MNN9, ROX1, DOS2, YJL064W, GFP) (Supplementary Fig. 1c).

Episomal plasmid BYP9686 (National Bio-Resource Project, Japan) was used for expressing GFP.

**Supplementary Table 1 Primer sequence used in this study**

| Primers | Sequence (5'-3') |
|---|---|
| tPG-E(F)_ASS | GCTAGCCGATATCCCTCTGTAAATTGAATTGAATTGAAATCGATAGATC |
| pPG-YB(R)_ASS | CGAGAGCCGTCATTGTTTTATATTTGTTGTAAAAAGTAG |
| YB-pPG(F)_ASS | ACAACAAATATAAAACAATGACGGCTCTCGCATATTACCAG |
| tAG-tHX(R)_ASS | GGACGGAGACAAAGAAACTTTAATTTGATTATGTTCTTTCTATTTGAATG |
| tHX-tAG(F)_ASS | TAGAAAGAACATAATCAAATTAAAGTTTCTTTGTCTCCGTCCCAC |
| tHX-I(R)_ASS | CGTTGGTGTTCTTGCTTTCTAATTTGCGAACACTTTTATTAATTCATG |
| I-tHX(F)_ASS | GAATTAATAAAAGTGTTCGCAAATTAGAAAGCAAGAACACCAACGGATC |
| I-pHX(R)_ASS | TTTTTTTAATTTTAATCAAAAAATGGGAAAAGAACAAGATCAGGATAAAC |
| pHX(F)_ASS | CTGATCTTGTTCTTTTCCCATTTTTTGATTAAAATTAAAAAAACTTTTTG |
| pHX-pTE(R)_ASS | GCTATGGTGTGTGGAATAGTACTCTCATCGCTAAGATCATTTG |
| pTE-pHX(F)_ASS | TAGCGATGAGAGTACTATTCCACACACCATAGCTTCAAAATGTTTC |
| pTE-E(R)_ASS | GTGAGGATGTTCGCGTAATCCATTTTGTAATTAAAACTTAGATTAGATTG |
| E-pTE(F)_ASS | ATCTAATCTAAGTTTTAATTACAAAATGGATTACGCGAACATCCTCACAG |
| E-tPG(R)_ASS | ATCGATTTCAATTCAATTCAATTTACAGAGGGATATCGGCTAGCTTTTTC |
| PRS404-AGO (F) | AAAGATAATATAGAAACAAAATGTCATCCAATTCGGAGGAGAAC |
| PRS404-DCR (R) | AATTGGGTACCGGGCCCCCCCTCGAGTCATTTTAAATTTTTTTTTATTCCTCATCGAACG |
| pHpGAP (F) | TCATCACTTATTTGCTCCCCAACCTGTCAAATACCTGCGTAAGGAATGCTGACTCTCTCC |
| pHpGAP (R) | TCCTCCGAATTGGATGACATTTTGTTTCTATATTATCTTTGTAC |
| AGO1_term (F) | TCATCACTTATTTGCTCCCCAACCTGTCAAATACCTGCGTAAGGAATGCTGACTCTCTCC |
| AGO1_term (R) | AATTGGGTACCGGGCCCCCCCTCGAGTCATTTTAAATTTTTTTTTATTCCTCATCGAACG |
| pPpTEF (F) | CTTCTTGGAGAGAGTCAGCATTCCTTACGCAGGTATTTGACAGGTTGGGGAGCAAATAAG |
| pPpTEF (R) | GCGCTTTTTTCTCTATTCATGTTGGCGAATAACTAAAATGTATGTAGTGAG |
| DCR1_term (F) | CATTTTAGTTATTCGCCAACATGAATAGAGAAAAAAGCGCCGATCTAAG |
| DCR1_term (R) | AAAAGCTGGAGCTCCACCGCGGTGGCGGCCGCTGGGGGGCTAGGCAAGTTATGTTTGAAG |
| 2mic(F)-EAD-GRC | CGAATTGGGTACCGGGCCCCCCCTCATATTTCCACGGACTATAGACTATAC |
| 2mic(R)-EAD-GRC | TCGATGAGGAATAAAAAAAAATTTAAAATGACTCGATATATCTGAACCAGTCCTAAAACG |
| FPLib (F) | ATTTCTATTCCAACAGAGCTCCACCGCGGTGGCGGCCGCGAGTCGACATCGAAGCGCGCC |
| FPLib (R) | CTTGTCAAAGGGTATTCCTGCAGGATGACCATGAAAGATGATTTTGTTATTTTAATTGTA |
| RPLib (F) | CATCTTTCATGGTCATCCTGCAGGAATACCCTTTGACAAGATTTTGTTATTTTAATTGTA |
| RPLib (R) | CGAGGTCGACGGTATCGATAAGCTTGATATCGCGGCCGCGAGTCGACATCGAAGCGCGCC |
| DPP1Mid600(F) | AATTAAAATAACAAAATCATCTTTCATGGTCATTACATTAACGATCTCACTATATC |
| DPP1Mid600(R) | AATTAAAATAACAAAATCTTGTCAAAGGGTATTTCTTCTGTAGAAAAAGTGTGCC |
| LPP1Mid600(F) | AATTAAAATAACAAAATCATCTTTCATGGTCATTGGTACCTAGGGGCCAAAACATC |
| LPP1Mid600(R) | AATTAAAATAACAAAATCTTGTCAAAGGGTATTACCAATGATGTCTGTGATCGATA |
| PAH1Mid600(F) | AATTAAAATAACAAAATCATCTTTCATGGTCATCTAGCAAGAAAGGAACAGCAGG |
| PAH1Mid600(R) | AATTAAAATAACAAAATCTTGTCAAAGGGTATTTCCTATTACCAAAGCCGGCAAAAA |
| ERG9Mid600(F) | AATTAAAATAACAAAATCATCTTTCATGGTCATTGACAGATTTCGAATCGATTC |

| | |
|---|---|
| ERG9Mid600(R) | AATTAAAATAACAAAATCTTGTCAAAGGGTATTTCTTTACATTGCCATGTAGCAC |
| EXG1Mid600(F) | AATTAAAATAACAAAATCATCTTTCATGGTCATAAGGTTTCAACCTTGTCAGAATTC |
| EXG1Mid600(R) | AATTAAAATAACAAAATCTTGTCAAAGGGTATTTCAAAACTCCGGTACCCCATTC |
| FLD1Mid600(F) | AATTAAAATAACAAAATCATCTTTCATGGTCATCTATTACCTGCCGATTCCTCTAAC |
| FLD1Mid600(R) | AATTAAAATAACAAAATCTTGTCAAAGGGTATTTTTTCGAAGCATCAAGTTTCTT |
| MNN9Mid600(F) | AATTAAAATAACAAAATCATCTTTCATGGTCATTATGATTTGAACAAATTGCACTC |
| MNN9Mid600(R) | AATTAAAATAACAAAATCTTGTCAAAGGGTATTACCCATCTGAGAGGCTATTTC |
| ROX1Mid600(F) | AATTAAAATAACAAAATCATCTTTCATGGTCATAGCCGGTAAGAAAGTCTAAGAAG |
| ROX1Mid600(R) | AATTAAAATAACAAAATCTTGTCAAAGGGTATTTTGAATTGTTGATACTGTTTAATC |
| DOS2Mid600(F) | AATTAAAATAACAAAATCATCTTTCATGGTCATACGATATGAAAAGACTACAAGTG |
| DOS2Mid600(R) | AATTAAAATAACAAAATCTTGTCAAAGGGTATTTCTTCGTCATCCCACTCAACTTC |
| YJL064W(F) | AATTAAAATAACAAAATCATCTTTCATGGTCATATGACACTTGTAGTATATCTAAC |
| YJL064W(R) | AATTAAAATAACAAAATCTTGTCAAAGGGTATTTCAGGCTAACACAATGAACAAC |
| GFPMid600 (F) | AATTAAAATAACAAAATCATCTTTCATGGTCATATTAGATGGTGATGTTAATG |
| GFPMid600 (R) | AATTAAAATAACAAAATCTTGTCAAAGGGTATTTGGTCTCTCTTTTCGTTG |
| DPP1-35(F) | ACATAGGGGCGAAATGGAGA |
| DPP1-183(R) | ACGTTCAGTTGTCGCATAAGG |
| LPP1-6(F) | CTCTGTCATGGCGGATGAGA |
| LPP1-136(R) | CGATGTTTTGGCCCCTAGGT |
| PAH1-932(F) | CTGGTTCCGATACTGAGGACG |
| PAH1-1020(R) | GCTCCCTGCTGTTCCTTTCT |
| ERG9-342(F) | CCCCGATGTGAAGGACAGAG |
| ERG9-471(R) | GTCGGCCATACCATTACCCA |
| EXG1-311(F) | GCCGTTTACAGAGCCATTGG |
| EXG1-419(R) | TGGAAAGCCCAGTAACCGAT |
| FLD1-18(F) | CCGTCCATTACAGTTTTTACAATGG |
| FLD1-163(R) | GGGGGACTACGTTAGAGGAATC |
| MNN9-233(F) | AAAGCGGCTGGTTGTTCAAC |
| MNN9-338(R) | TTGACAGCTGCTTCTGACGT |
| ROX1-171(F) | ACCGGAAGATAAGGCACACTG |
| ROX1-313(R) | GCTGCTGTTGCTCGATTTCC |
| DOS2-122(F) | ACAAAACAAATGAGGCCTTCCA |
| DOS2-270(R) | TGCAGTCTCTGTAGTTTCATTGC |
| YJL064W-78(F) | CAAAAACTGCAGATGGCCGT |
| YJL064W-187(R) | AGCAGCAGCAACAACAACAG |
| PDA1-497(F) | CTTATGGTAAGGGTGGTTCCATG |
| PDA1-606(R) | TGGTGAGCAAAAGCTAAACCTG |

**Supplementary Table 2 Primers for plasmid construction**

| Gene fragment | Template DNA | Primer (F) | Primer (R) |
|---|---|---|---|
| A1 | pEU20PG | tPG-E(F)_ASS | pPG-YB(R)_ASS |
| A2 | pδL PGxdCrtYB | YB-pPG(F)_ASS | tAG-tHX(R)_ASS |
| A3 | S288C genome | tHX-tAG(F)_ASS | tHX-I(R)_ASS |
| A4 | pδL PGxdCrtI | I-tHX(F)_ASS | I-pHX(R)_ASS |
| A5 | *S. cerevisiae* YPH499 genome | pHX(F)_ASS | pHX-pTE(R)_ASS |
| A6 | *S. cerevisiae* YPH499 genome | pTE-pHX(F)_ASS | pTE-E(R)_ASS |
| A7 | pδL PGxdCrtE | E-pTE(F)_ASS | E-tPG(R)_ASS |
| B1 | pRS404 | PRS404-AGO (F) | PRS404-DCR (R) |
| B2 | *H. polymorpha* NBRC1476 genome | pHpGAP (F) | pHpGAP (R) |
| B3 | *S. castellii* NBRC1992 genome | AGO1_term (F) | AGO1_term (R) |
| B4 | *P. pastoris* GS115 genome | pPpTEF (F) | pPpTEF (R) |
| B5 | *S. castellii* NBRC1992 genome | DCR1_term (F) | DCR1_term (R) |
| B6 | pEUPGGFP | 2mic(F)-EAD-GRC | 2mic(R)-EAD-GRC |
| C1 | pδU_LibDLDH | FPLib (F) | FPLib (R) |
| C2 | pδU_LibDLDH | RPLib (F) | RPLib (R) |
| C3 | *S. cerevisiae* YPH499 genome | DPP1Mid600(F) | DPP1Mid600(R) |
| C4 | *S. cerevisiae* YPH499 genome | LPP1Mid600(F) | LPP1Mid600(R) |
| C5 | *S. cerevisiae* YPH499 genome | PAH1Mid600(F) | PAH1Mid600(R) |
| C6 | *S. cerevisiae* YPH499 genome | ERG9Mid600(F) | ERG9Mid600(R) |
| C7 | *S. cerevisiae* YPH499 genome | EXG1Mid600(F) | EXG1Mid600(R) |
| C8 | *S. cerevisiae* YPH499 genome | FLD1Mid600(F) | FLD1Mid600(R) |
| C9 | *S. cerevisiae* YPH499 genome | MNN9Mid600(F) | MNN9Mid600(R) |
| C10 | *S. cerevisiae* YPH499 genome | ROX1Mid600(F) | ROX1Mid600(R) |
| C11 | *S. cerevisiae* YPH499 genome | DOS2Mid600(F) | DOS2Mid600(R) |
| C12 | *S. cerevisiae* YPH499 genome | YJL064W(F) | YJL064W(R) |
| C13 | BYP9686 | GFPMid600 (F) | GFPMid600 (R) |

### Supplementary Table 3 Gene sequence for RNA interference

| Gene | Sequence for double stranded RNA |
|---|---|
| *DPP1* | TACATTAACGATCTCACTATATCGCATCCTTATGCGACAACTGAACGTGTAAATAACAACATGTTGTTTGTTTATAGT<br>TTTGTCGTGCCATCTTTAACCATATTGATAATTGGTTCCATTTTGGCCGATAGAAGACATTTGATTTTTATTTTGTACA<br>CATCTCTCCTTGGTTTATCACTCGCTTGGTTCAGTACGAGTTTCTTTACAAACTTCATCAAGAATTGGATTGGAAGACT<br>AAGACCAGATTTTCTAGATCGTTGCCAACCTGTTGAAGGCTTGCCATTGGACACTTTATTTACTGCAAAAGATGTGTG<br>TACGACTAAGAATCACGAACGTCTGTTGGATGGGTTTAGGACAACTCCGTCAGGTCATTCAAGTGAAAGCTTTGCAG<br>GACTGGGTTATTTGTACTTCTGGCTATGTGGGCAACTTTTGACTGAATCACCGTTGATGCCTTTATGGAGAAAAATGG<br>TGGCCTTTCTACCACTGTTAGGAGCTGCACTAATTGCTCTATCCAGAACTCAAGATTACAGACATCATTTCGTCGATG<br>TAATTTTAGGGTCTATGTTGGGTTATATAATGGCACACTTTTTCTACAGAAGA |
| *LPP1* | TGGTACCTAGGGGCCAAAACATCGAATTTAGTCTTGATGACCCCAGTATATCAAAACGTTATGTACCTAACGAACTC<br>GTGGGCCCACTAGAATGTTTGATTTTGAGTGTTGGACTGAGTAACATGGTCGTCTTCTGGACCTGCATGTTTGACAAG<br>GACTTACTGAAGAAGAATAGAGTAAAGAGACTAAGAGAGAGGCCGGACGGAATCTCGAACGATTTTCACTTCATGC<br>ATACTAGCATTCTATGTCTGATGCTGATTATAAGCATAAATGCTGCCCTAACAGGCGCCTTAAAGTTGATTATAGGAA<br>ACTTGAGGCCTGACTTTGTTGATAGATGTATACCTGACCTCCAAAAGATGAGTGATTCAGATTCTTTGGTTTTTGGCT<br>TGGACATTTGCAAGCAGACTAACAAATGGATTCTATACGAAGGCTTAAAAAGCACTCCAAGCGGACATTCAAGTTTC<br>ATAGTCAGTACCATGGGCTTTACATATCTTTGGCAAAGGGTTTTCACCACACGCAATACAAGAAGTTGCATTTGGTGC<br>CCTTTATTAGCTCTAGTAGTAATGGTTTCAAGGGTTATCGATCACAGACATCATTGGT |
| *PAH1* | CTAGCAAGAAAGGAACAGCAGGGAGCGGTGAGACCGAGAAAAGATACATACGAACGATAAGATTGACTAATGACC<br>AGTTAAAGTGCCTAAATTTAACTTATGGTGAAAATGATCTGAAATTTTCCGTAGATCACGGAAAAGCTATTGTTACGT<br>CAAAATTATTCGTTTGGAGGTGGGATGTTCCAATTGTTATCAGTGATATTGATGGCACCATCACAAAATCGGACGCTT<br>TAGGCCATGTTCTGGCAATGATAGGAAAAGACTGGACGCACTTGGGTGTAGCCAAGTTATTTAGCGAGATCTCCAGG<br>AATGGCTATAATATACTCTATCTAACTGCAAGAAGTGCTGGACAAGCTGATTCCACGAGGAGTTATTTGCGATCAAT<br>TGAACAGAATGGCAGCAAACTACCAAATGGGCCTGTGATTTTATCACCCGATAGAACGATGGCTGCGTTAAGGCGGG<br>AAGTAATACTAAAAAAACCTGAAGTCTTTAAAATCGCGTGTCTAAACGACATAAGATCCTTGTATTTTGAAGACAGT<br>GATAACGAAGTGGATACAGAGGAAAAATCAACACCATTTTTTGCCGGCTTTGGTAATAGGA |
| *ERG9* | TGACAGATTTCGAATCGATTCTTATTGAATTCCACAAATTGAAACCAGAATATCAAGAAGTCATCAAGGAGATCACC<br>GAGAAAATGGGTAATGGTATGGCCGACTACATCTTAGATGAAAATTACAACTTGAATGGGTTGCAAACCGTCCACGA<br>CTACGACGTGTACTGTCACTACGTAGCTGGTTTGGTCGGTGATGGTTTGACCCGTTTGATTGTCATTGCCAAGTTTGC<br>CAACGAATCTTTGTATTCTAATGAGCAATTGTATGAAAGCATGGGTCTTTTCCTACAAAAAACCAACATCATCAGAG<br>ATTACAATGAAGATTTGGTCGATGGTAGATCCTTCTGGCCCAAGGAAATCTGGTCACAATACGCTCCTCAGTTGAAG<br>GACTTCATGAAACCTGAAAACGAACAACTGGGGTTGGACTGTATAAACCACCTCGTCTTAAACGCATTGAGTCATGT<br>TATCGATGTGTTGACTTATTTGGCCGGTATCCACGAGCAATCCACTTTCCAATTTTGTGCCATTCCCCAAGTTATGGCC<br>ATTGCAACCTTGGCTTTGGTATTCAACAACCGTGAAGTGCTACATGGCAATGTAAAGA |
| *EXG1* | AAGGTTTCAACCTTGTCAGAATTCCTATCGGTTACTGGGCTTTCCAAACTTTGGACGATGATCCTTATGTTAGCGGCC<br>TACAGGAATCTTACCTAGACCAAGCCATCGGTTGGGCTAGAAACAACAGCTTGAAAGTTTGGGTTGATTTGCATGGT<br>GCCGCTGGTTCGCAGAACGGGTTTGATAACTCTGGTTTGAGAGATTCATACAAGTTTTTGGAAGACAGCAATTTGGC<br>CGTTACTACAAATGTCTTGAACTACATATTGAAAAAATACTCTGCGGAGGAATACTTGGACACTGTTATTGGTATCGA<br>ATTGATTAATGAGCCATTGGGTCCTGTTCTAGACATGGATAAAATGAAGAATGACTACTTGGCACCTGCTTACGAAT<br>ACTTGAGAAACAACATCAAGAGTGACCAAGTTATCATCATCCATGACGCTTTCCAACCATACAATTATTGGGATGAC<br>TTCATGACTGAAAACGATGGCTACTGGGGTGTCACTATCGACCATCATCACTACCAAGTCTTTGCTTCTGATCAATTG<br>GAAAGATCCATTGATGAACATATTAAAGTAGCTTGTGAATGGGGTACCGGAGTTTTGA |
| *FLD1* | CTATTACCTGCCGATTCCTCTAACGTAGTCCCCCTTAATACGTTCAACATTTTAAATGGCGTACAATTTGGTACAAAA<br>TTCTTCCAATCTATTAAAAGCATTCCGGTAGGTACAGATCTGCCGCAAACAATAGACAATGGCTTATCACAGTTAATC<br>CCCATGCGTGACAACATGGAATACAAGCTCGATCTAAACCTACAGCTTTATTGCCAGAGCAAAACTGACCATTTAAA<br>TTTAGACAATTTGTTAATTGATGTTTACAGAGGTCCAGGCCCGCTATTGGGTGCTCCTGGAGGAAGTAACAGCAAAG<br>ATGAAAAAATCTTTCACACTTCTAGACCTATTGTCTGCCTCGCACTGACGGATTCCATGTCGCCTCAGGAGATCGAAC<br>AACTAGGCCCATCACGTCTAGACGTTTACGATGAAGAATGGCTAAATACAATAAGAATAGAGGACAAAATATCGTT<br>AGAGTCTTCATATGAAACAATCTCGGTGTTCTTGAAAACGGAGATTGCCCAAAGAAATCTAATAATACATCCAGAAA<br>GTGGGATTAAGTTTAGGATGAATTTTGAGCAGGGATTAAGAAACTTGATGCTTCGAAAA |

| | |
|---|---|
| *MNN9* | TATGATTTGAACAAATTGCACTCTACGTCAGAAGCAGCTGTCAATAAGGAGCATATTTTGATATTGACTCCAATGCA<br>AACATTTCATCAACAATACTGGGACAATTTGCTGCAGCTAAATTACCCTCGGGAATTGATTGAATTGGGGTTCATTAC<br>ACCAAGAACAGCCACTGGTGACTTGGCCTTAAAGAAATTGGAGAATGCTATTAAAAAGGTTCAAACGGACAAGAAA<br>ACTCAAAGATTTAGTAAAATTACTATTTTGCGACAGAATTCCCAGAGTTTTGATAAGTTGATGGAGAAGGAAAGACA<br>CGCTTTAGATGTTCAAAAGGAAAGACGTGCAGCAATGGCTTTGGCGCGCAATGAATTACTATTCTCCACCATAGGAC<br>CTCACACTTCTTGGGTGCTGTGGCTAGATGCCGATATTATAGAGACACCACCATCTTTAATTCAAGACATGACCAAAC<br>ACAACAAAGCTATCTTAGCTGCAAACATTTATCAAAGATTTTACGATGAAGAGAAGAAGCAACCATCAATCAGACCA<br>TACGATTTCAACAACTGGCAAGAAAGTGACACCGGTTTAGAAATAGCCTCTCAGATGGGT |
| *ROX1* | AGCCGGTAAGAAAGTCTAAGAAGAAGCAACTACTTTTGAAGGAAATCGAGCAACAGCAGCAGCAACAACAGAAAG<br>AACAGCAGCAGCAGAAACAGTCACAACCGCAATTACAACAGCCCTTTAACAACAATATAGTTCTTATGAAAAGAGC<br>ACATTCTCTTTCACCATCTTCCTCGGTGTCAAGCTCGAACAGCTATCAGTTCCAATTGAACAATGATCTTAAGAGGTT<br>GCCTATTCCTTCTGTTAATACTTCTAACTATATGGTCTCCAGATCTTTAAGTGGACTACCTTTGACGCATGATAAGACG<br>GCAAGAGACCTACCACAGCTGTCATCTCAACTAAATTCTATTCCATATTACTCAGCTCCACACGACCCTTCAACGAGA<br>CATCATTACCTCAACGTCGCTCAAGCTCAACCAAGGGCTAACTCGACCCCTCAATTGCCCTTTATTTCATCCATTATC<br>AACAACAGCAGTCAAACACCGGTAACTACAACTACCACATCCACAACAACTGCGACATCTTCTCCTGGGAAATTCTC<br>CTCTTCTCCGAACTCCTCTGTACTGGAGAACAACAGATTAAACAGTATCAACAATTCAA |
| *DOS2* | ACGATATGAAAAGACTACAAGTGCATTCAAAAAGTTAGTAATCGAGAAAGATGACGGAATTGAAATTAACTTGCCA<br>ATTAGCAATGAAACTACAGAGACTGCACAAAAGTATTTGAAGAAACTAGACGAGAATATTCACAGCGTGGAAAGTC<br>TAGCCCAGTCATATTGGAGCAAAATGAAAACTAAGAATTTTTGGTCTGGCTTCAGTAGCTTCGATAATGCTGCAGAA<br>AATGACTCTAATGACAAGGATGAGAATTCGAAAGAAAATGAAATTGCTGTGGGTGGAAATAGAACAGAAGCCGAAC<br>TAAGGACATTATCTAAAGATAAATCGGTTTATTTAGACAATAAAATGGATTTGCAACTGGACCCGTTTGACGTGGAT<br>GAAAAAACTGAGGAGATATGTTCTATTTTACAGGGCGACAAAGATATCTCCAAATTAATGAACGACATCGTACCCCA<br>TAAAATCAGCTATAAAGATTTCTGGCACATCTATTTCTTACAAAGAAACAAAATTCTAGATAAAGAAAGTAAAAGGA<br>AAGAAATATTGTCCAAAAAGGAAAAGGAAACGGAGGAAAAAGAAGTTGAGTGGGATGACGAAGA |
| *YJL064W* | ATGACACTTGTAGTATATCTAACTCGGTTTTCTTCCACTAGAAGCCTCAAATTCTTTGTTGTTGGTATCACATCATTCA<br>AAAACTGCAGATGGCCGTCAGAAGAGTGCACAATTGCGGCAGAGATGTCATCGTACGACAGCGTGAGTTCATCTGG<br>GAGCGGCGGTACCTGTTGTTGTTGCTGCTGCTGTTGCCTATGTAGGGACTCATGTGTCAGCACCTGGACGAAGAATTC<br>TGTAGCAAACGCTGTGGCTACGAATGCGTCATCCGAAGTGTCCATATATTCTGGATCATTCTTGGCTATTCTCTGCAC<br>TTTTTCTACAGGTAACTTGGGCGAACACAGAGGTGCAGACGCCGTTTCTTTACCACTAGTCTCGTTGTTCATTGTGTT<br>AGCCTGA |
| *GFP* | ATTAGATGGTGATGTTAATGGGCACAAATTTTCTGTCAGTGGAGAGGGTGAAGGTGATGCAACATACGGAAAACTTA<br>CCCTTAAATTTATTTGCACTACTGGAAAACTACCTGTTCCATGGCCAACACTTGTCACTACTTTCACTTATGGTGTTCA<br>ATGCTTTTCAAGATACCCAGATCATATGAAACGGCATGACTTTTTCAAGAGTGCCATGCCCGAAGGTTATGTACAGG<br>AAAGAACTATATTTTTCAAAGATGACGGGAACTACAAGACACGTGCTGAAGTCAAGTTTGAAGGTGATACCCTTGTT<br>AATAGAATCGAGTTAAAAGGTATTGATTTTAAAGAAGATGGAAACATTCTTGGACACAAATTGGAATACAACTATAA<br>CTCACACAATGTATACATCATGGCAGACAAACAAAAGAATGGAATCAAAGTTAACTTCAAAATTAGACACAACATT<br>GAAGATGGAAGCGTTCAACTAGCAGACCATTATCAACAAAATACTCCAATTGGCGATGGCCCTGTCCTTTTACCAGA<br>CAACCATTACCTGTCCACACAATCTGCCCTTTCGAAAGATCCCAACGAAAAGAGAGACCA |

**Supplementary Fig. 1 Plasmids used in this study**

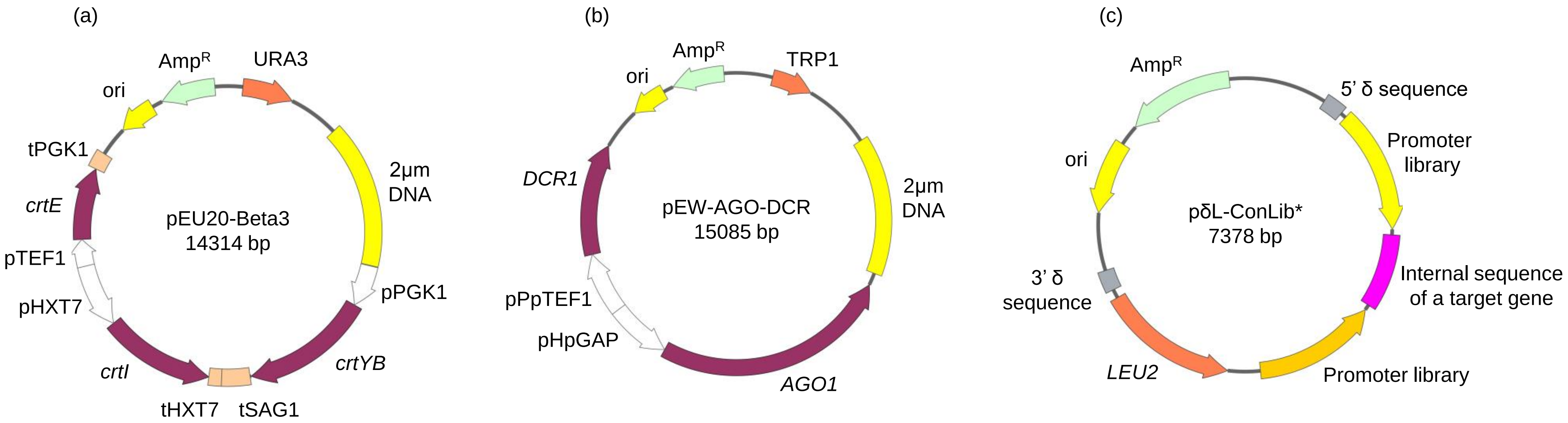


Plasmid for (a) carotenoid production and (b) expression of Argonaute and Dicer and (c) plasmid library for double-stranded RNA expression.

**Supplementary Fig. 2 Evaluation of RNA interference ability using GFP**

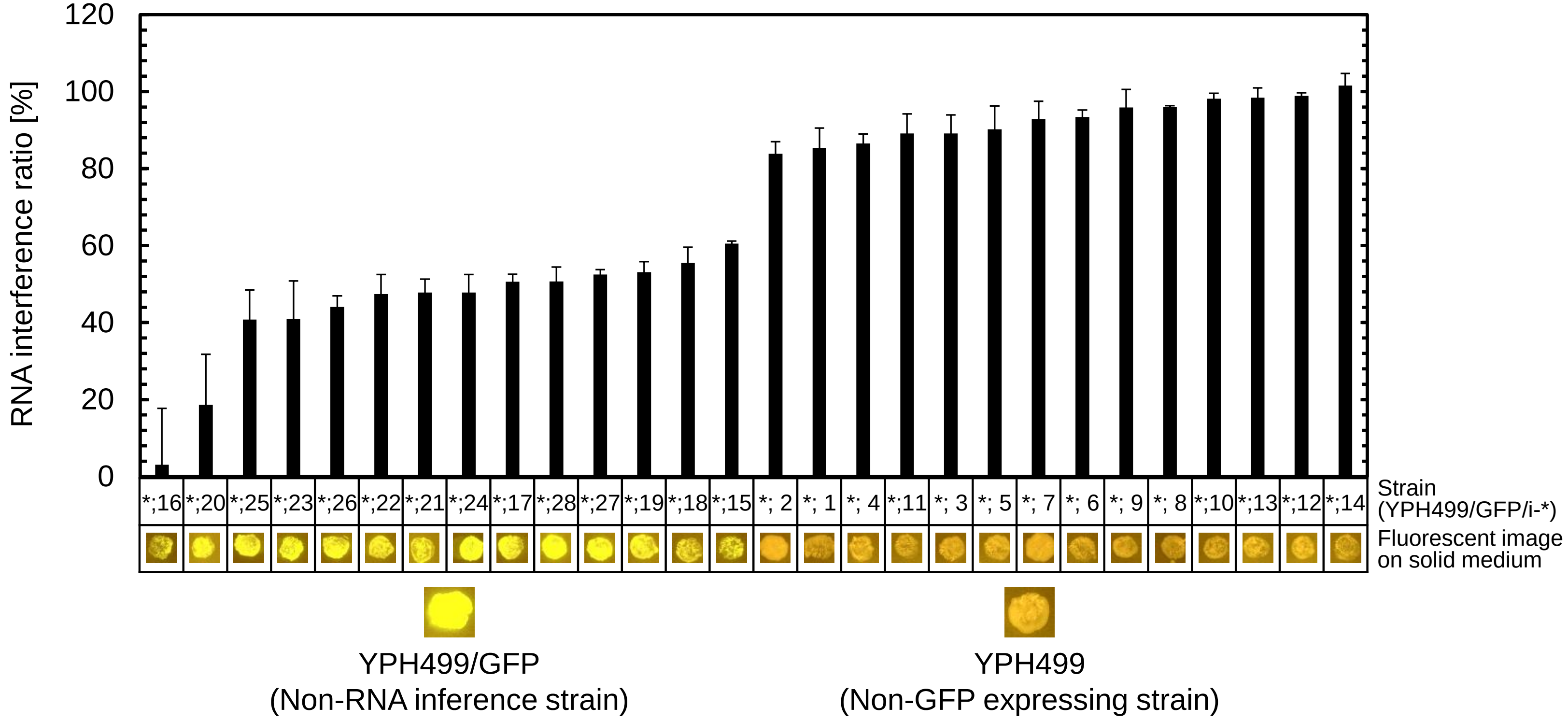


RNA interference ability was evaluated using the GFP expression system. A yeast strain YPH499 expressing GFP, Argonaute, and Dicer, as well as double-stranded RNA that interfere with GFP expression at various intensities, was constructed and designated YPH499/GFP/i-* (*; 1-28). The fluorescence intensity of the constructed yeast was measured after 48 h of cultivation. The measured fluorescence intensities are shown in ascending order from left to right as the RNA interference ratio (Equation S1). Data are average of 3 independent experiments and error bar represents standard deviation. The constructed GFP interference strains showed diverse fluorescence intensities, confirming that the constructed RNA interference technology can be used to suppress GFP expression at various intensities.

$$\text{RNA interference ratio [\%]} = \left[1 - \frac{\text{Fluorescence value of GFP w/ RNA interference}}{\text{Fluorescence value of GFP w/o RNA interference}}\right] \times 100 \qquad \text{(Equation S1)}$$

**Supplementary Fig. 3 Evaluation of RNA interference stability using GFP**

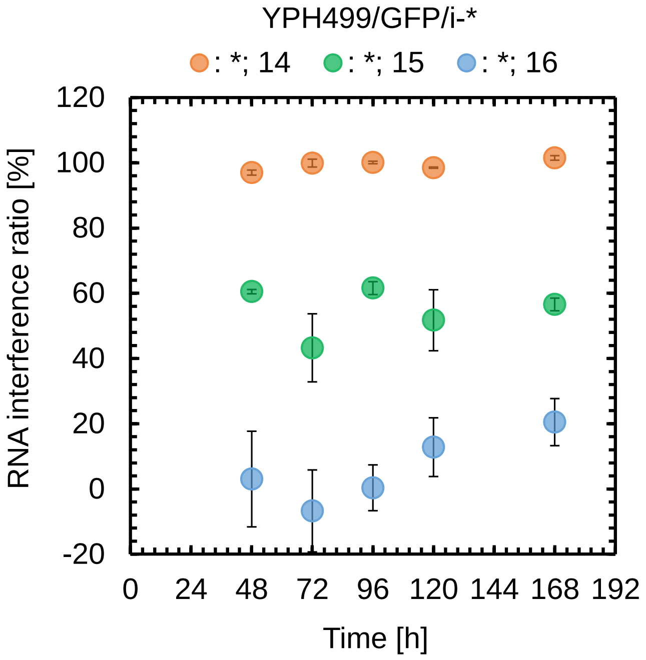


RNA interference stability was also evaluated using the GFP expression system. In the results shown in Supplementary Fig. 1, YPH499/GFP/i-14, YPH499/GFP/i-15, and YPH499/GFP/i-16, which showed high, intermediate, and low RNA interference ratio, were cultured and their time course of RNA interference ratio were evaluated. Data are average of 3 independent experiments and error bar represents standard deviation. All strains showed a nearly constant RNA interference ratio up to 168 h of culture. Therefore, it was confirmed that the constructed RNA interference system can stably maintain RNA interference ability over a long period of time.

**Reference**

Ma, H.; Kunes, S.; Schatz, P. J.; Botstein, D. Plasmid construction by homologous recombination in yeast. *Gene* **1987**, *58*, 201-216.

Yamada, R.; Wakita, K.; Mitsui, R.; Nishikawa, R.; Ogino, H. Efficient production of 2,3-butanediol by recombinant *Saccharomyces cerevisiae* through modulation of gene expression by cocktail delta-integration. *Bioresour. Technol.* **2017**, *245*, 1558-1566.
